# DARKIN: A zero-shot benchmark for phosphosite–dark kinase association using protein language models

**DOI:** 10.1101/2025.08.27.672558

**Authors:** Emine Ayşe Sunar, Zeynep Işık, Mert Pekey, Ramazan Gökberk Cinbiş, Oznur Tastan

## Abstract

**Motivation:** Protein Language Models (pLMs) have emerged as powerful tools for capturing the intricate information encoded in protein sequences, facilitating various downstream protein prediction tasks. With numerous pLMs available, there is a critical need for diverse benchmarks to systematically evaluate their performance across biologically relevant tasks. Here, we introduce DARKIN, a zero-shot classification benchmark designed to assign phosphosites to understudied kinases, termed dark kinases. Kinases, which catalyze phosphorylation, are central to cellular signaling pathways. While phosphoproteomics enables the large-scale identification of phosphosites, determining the cognate kinase responsible for the phosphorylation event remains an experimental challenge.

**Results:** In DARKIN, we prepared training, validation, and test folds that respect the zero-shot nature of this classification problem, incorporating stratification based on kinase groups and sequence similarity. We evaluated multiple pLMs using two zero-shot classifiers: a novel, training-free k-NN-based method, and a bilinear classifier. Our findings indicate that ESM, ProtT5-XL, and SaProt exhibit superior performance on this task. DARKIN provides a challenging benchmark for assessing pLM efficacy and fosters deeper exploration of under-characterized (dark) kinases by offering a biologically relevant test bed.

**Implementation:** The DARKIN benchmark data and the scripts for generating additional splits are publicly available at: https://github.com/tastanlab/darkin

**Supplementary information:** Supplementary data are available at *Bioinformatics* online.

## Introduction

Building on the success of large language models (LLMs) in natural language processing [Zhao et al., 2023], protein language models (pLMs) have been developed to capture the complex information embedded within protein sequences [Rao et al., 2019, Elnaggar et al., 2021, Lin et al., 2023, Meier et al., 2021, Lin et al., 2022, Brandes et al., 2022, Ferruz et al., 2022, Geffen et al., 2022, Elnaggar et al., 2023, Su et al., 2023, Zhang et al., 2025, Hayes et al., 2025, ESM Team, 2024, Fournier et al., 2024, Peng et al., 2025, Wang et al., 2024, Ouyang-Zhang et al., 2024]. By generating semantic representations of proteins, pLMs enable a broad range of sequence-based prediction tasks. However, as more pLMs become available, systematically benchmarking their performance is essential to determine their reliability and applicability in diverse biological contexts. Previous work has compared the pLMs in their ability to predict proteins’ functional properties [Unsal et al., 2022, Schmirler et al., 2024, Zhang et al., 2025], and functional motifs [SAV, 2023]. In this work, we provide a novel biologically relevant zero-shot prediction benchmark for phosphosite-dark kinase associations and compare the pLMs in terms of their ability to capture intrinsic sequence properties within this challenging task.

Phosphorylation events are key regulators of protein function in signal transduction pathways, and their dysfunction is associated with many diseases [Gaestel et al., 2009, Wu et al., 2023, Müller et al., 2015]. Kinases are the enzymes that catalyze the phosphorylation of other proteins in a target-specific manner [Hunter, 1995]. For this reason, kinases are major drug targets in diseases such as cancer, infectious diseases, and neurological disorders [Blume-Jensen and Hunter, 2001, Cohen et al., 2021]. Phosphorylation involves transferring a phosphate from adenosine 5’-triphosphate (ATP) to amino acid residues [Cohen, 2002]. These phosphorylated residues, referred to as *phosphosites*, are integral to modulating the protein’s structure and function.

Although high-throughput phosphoproteomics enables the identification of phosphosites at the proteome level, experimentally determining the kinase responsible for a phosphorylation event remains a major challenge. Notably, more than 95% of reported human phosphosites have no known cognate kinase [Needham et al., 2019], and 25% of the kinases are yet to be assigned to a phosphorylation event; for about 35% of the kinases, there are 1-10 phosphosites have been identified (Supplementary Figure S1). Consequently, most of the phosphoproteome and the kinome are in the dark [Needham et al., 2019, Vella et al., 2022, Deznabi et al., 2020]. Associating ‘orphan’ phosphosites to their respective kinases is an important task that would help understand the biological function of these phosphorylation events and discover new drug targets [Berginski et al., 2021, Deznabi et al., 2020, Needham et al., 2019]. In this work, given a phosphosite, we aim to predict the dark kinase associated with this phosphosite.

The contributions of this work can be summarized as follows: (i) We present a reproducible benchmark dataset for predicting dark kinase-phosphosite associations. The task is formulated as given a phosphosite, predict the associated dark kinase of that site. (ii) We propose a strategy to split the dataset into train, validation, and test splits for this zero-shot multi-class prediction task, with stratification based on kinase groups, the number of phosphosites per kinase, and kinase sequence similarity. (iii) We present a novel, training-free, k-NN-based zero-shot classification method for assessing the performance of protein language models (pLMs) under the task of predicting the dark kinase of a given phosphosite. (iv) We evaluate and compare various pLMs using two distinct zero-shot classification approaches.

## Methods

### Problem Description

Let 𝒳 denote the space of phosphosite sequences and 𝒴 denote the set of all human kinases. The task of kinase-phosphosite association prediction involves identifying the kinase *y* ∈ 𝒴 most likely to catalyze the phosphorylation of a given phosphosite sequence *x* ∈ 𝒳. Since a phosphosite can be phosphorylated by multiple kinases, we frame the problem as a multilabel classification task. We denote the training kinases as 𝒴_*tr*_ ⊂ 𝒴 and the test kinases as 𝒴_*te*_ ⊂ 𝒴. The set 𝒴_*te*_ comprises the zero-shot classes, and the training and test kinase sets are disjoint. The training data, *D*_*tr*_ = (*x*_*i*_, *y*_*i*_), *i* = 1, … , *N*_*tr*_, consists of pairings of train kinases with their associated phosphosites, where *y*_*i*_ ∈ 𝒴_*tr*_. Similarly, the test data contains phosphosite pairings of the test kinases 𝒴_*te*_.

### Dataset Curation and Processing

The DARKIN dataset is built on human kinases and their associated phosphosites. Several publicly available human kinase lists are available, yet they partially overlap due to ambiguities in defining kinase domains. The most widely used and oldest list is the 518 human kinase set defined by Manning et al. [2002]. Other sources, such as kinasecom (http://kinase.com/), Eid et al. [2017], Consortium [2023], and Moret et al. [2020], provide alternative kinase lists with some variations. For the current work, we resort to an up-to-date list from Moret et al. [2020], which includes 557 human kinases, each containing at least one kinase domain.

We obtained experimentally validated kinase-phosphosite associations from the PhosphoSitePlus [Hornbeck et al., 2012] (downloaded in May 2023). Kinase-phosphosite associations, which are related to non-human kinases, are removed. We did not apply the same restriction to substrates, as substrates from the model organisms are used to probe the interactions. We removed kinase isoforms and fusion kinases and used the canonical form specified in the UniProt human proteome [Bairoch et al., 2005] (downloaded May 2023). Phosphosites are represented as 15-residue amino acid sequences, including seven residues flanking the phosphosite on both sides. Previous work has shown that phosphosite sequences of length 15 or shorter led to better performances [Hornbeck et al., 2014, Trost and Kusalik, 2011, Wagih et al., 2015, Deznabi et al., 2020]. Padding was applied to ensure the phosphosite remains centered when it is near the N- or C-terminus of the protein.

Protein sequences were retrieved from UniProt via the API [Ahmad et al., 2025] (accessed December 2023). If the substrate could not be uniquely mapped to a Uniprot ID, we removed all phosphosite-kinase associations of these substrates. We retrieved the kinase domain sequences using the domain indices provided in Moret et al. [2020]. Kinases are categorized into groups and families by Manning et al. [2002] according to their domain sequence similarities. We retrieved the kinase family and group information from Moret et al. [2020]. Missing group and family information was imputed according to their similarity to other kinases. We defined a kinase group Other2 and a kinase family `otherFamily` for kinases that cannot be assigned to a family or group due to their dissimilarity to the rest of the groups. Another categorical information regarding kinases is the Enzyme Commission (EC) categorization. EC numbers categorize kinases according to their functionality. We downloaded Enzyme Commission (EC) numbers of the kinases (downloaded July 2023) [Bairoch, 2000]. We obtained protein structure data from the AlphaFold Protein Structure Database using AlphaFoldAPI at EBI (https://alphafold.ebi. ac.uk [Jumper et al., 2021, Varadi et al., 2022] and PDBe [Varadi et al., 2020]. For isoform proteins lacking structural data in AlphaFold and PDBe, we used ColabFold to predict 3D structures [Mirdita et al., 2022].

### Evaluated Protein Language Models and Baseline Encodings

We selected pLMs whose models were accessible, reported to perform well in the literature, and were recent. Table 1 presents the pLMs we evaluated, along with their key properties. For more efficient processing, we computed the column-wise average of the embedding for all pLMs, excluding the vectors corresponding to the padding (PAD) token. For pLMs with a classification (CLS) token, we used the embeddings corresponding to this token to summarize the overall representation.

**Table 1.** The protein Language Models (pLMs) compared in this study.

| PLM | Dataset | Vector Size | Model Size | Representation | Objective | Citation |
| --- | --- | --- | --- | --- | --- | --- |
| TAPE | PFAM | 768 | 38M | Sequence | Sequence-based, Structural Feature Prediction | Rao et al., 2019 |
| ProtBERT | BFD100, UniRef100 | 1024 | 420M | Sequence | Sequence-based, Structural, Physicochemical Feature Prediction | Elnaggar et al., 2021 |
| ProtALBERT | UniRef100 | 4096 | 224M |  |  |  |
| ProtT5-XL | BFD100 | 1024 | 3B |  |  |  |
| ESM1B | UniRef50 | 1280 | 650M | Sequence | Structural, Physicochemical Feature Prediction | Lin et al., 2023 |
| ESM1v | UniRef90 | 1280 | 650M | Sequence | Sequence Variant Prediction | Meier et al., 2021 |
| ESM2 | UniRef50 | 1280 | 650M | Sequence | Structural Feature, Contact Prediction | Lin et al., 2022 |
| ProteinBERT | UniRef90 | 1562 | 16M | Sequence | Sequence-based Feature, GO Annotation Prediction | Brandes et al., 2022 |
| ProtGPT2 | UniRef50 | 1280 | 738M | Subword | Protein Design and Engineering | Ferruz et al., 2022 |
| DistilProtBERT | UniRef50 | 1024 | 230M | Sequence | Sequence-based, Structural, Physicochemical Feature Prediction | Geffen et al., 2022 |
| Ankh | UniRef50 | 1536 | 1.5B | Sequence | General Purpose Modeling | Elnaggar et al., 2023 |
| SaProt | AlphaFold2, PDB | 1280 | 650M | Sequence, Structure | Structure-Aware Feature, Mutation Effect Prediction | Su et al., 2023 |
| ESM3 | UniRef, MGnify90, JGI, OAS PDB, InterPro, InterProScan | 1536 | 1.4B | Sequence, Structure, Function | Protein Generation | Hayes et al., 2025 |
| ESMC | UniRef, MGnify, JGI | 1152 | 600M | Sequence | Sequence-based Feature, Contact Prediction | ESM Team, 2024 |
| ISM2 | Uniclust30, PDB | 1280 | 650M | Sequence | Sequence-based, Structural, Functional Feature Prediction | Ouyang-Zhang et al., 2024 |
| DPLM | UniRef50 | 960 | 650M | Sequence | Conditional and Unconditional Protein Generation | Wang et al., 2024 |
| AMPLIFY | UniRef, OAS, SCOP, UniProt | 1280 | 350M | Sequence | Structural Feature, Contact Prediction | Fournier et al., 2024 |
| PTM-Mamba | UniProt Swiss-Prot PTM | 768 | 220M (Mamba) + 650M (ESM2)* | Sequence | PTM-related Prediction, PTM Discovery | Peng et al., 2025 |

\*This is the parameter size of ESM2, which is also used in PTM-Mamba. The versions of the models are specified in Supplementary Table S4.

In addition to the pLM, we used the following encodings as the baseline representations:

i. **One-hot encoding:** The input sequence is expressed as a binary vector of amino acids.
ii. **BLOSUM62 encoding:** The encoding uses the row corresponding to a particular amino acid in the BLOSUM62 matrix, which represents the probability of substitution of that amino acid by any other amino acid.
iii. **NLF encoding:** NLF captures the physicochemical properties of amino acids and is determined by a non-linear Fisher transform [Nanni and Lumini, 2011]. The representations are computed using the epitope prediction tool [Farrell, 2021].
iv. **ProtVec:** ProtVec is a skip-gram neural network model trained to provide a continuous representation of protein sequences [Asgari and Mofrad, 2015]. ProtVec provides a 100-dimensional embedding for each 3-gram, and the average embedding is used to represent the sequence.

### Evaluation Splits

In the zero-shot learning evaluation protocol, it is crucial to ensure class separation during model training and hyperparameter tuning [Xian et al., 2017]. Therefore, examples are divided into train, validation, and test sets based on their associated class labels. In our earlier work, DeepKinZero evaluation Deznabi et al. [2020], we partitioned the data into training, validation, and test sets according to the number of phosphosites associated with each kinase. Kinases with more than five phosphosites were assigned as training classes, while kinases associated with exactly five phosphorylation sites were designated as validation kinases. The remaining kinases, each with fewer than five phosphosites, form the test or zero-shot classes. Thus, in this setup, the zero-shot kinases represent the dark kinases, whereas the training classes are light kinases. While this splitting strategy closely mirrors the real-world scenario of the deployed model, the limited number of examples for each class in the test set complicates the reliable estimation of evaluation performance. Therefore, we establish a setup where a portion of the well-studied kinases (light kinases) are held out as zero-shot classes and are excluded from the training process. Thus, imitating that light kinases are dark kinases. We follow this strategy to ensure that we have enough data from each kinase in the test set to report a more robust evaluation of the performance metrics. When creating the splits, we consider the following criteria to ensure a fair evaluation of data splits:

i. **Number of phosphosites per kinase:** To ensure robust evaluation, we set a minimum threshold for the number of kinase-phosphosite pairs associated with each kinase in the test and validation sets. This prevents relying on very few data points related to a specific kinase class, minimizing inaccurate and unstable results. Thus, we invert the roles of light and dark kinases in evaluation: the test data include well-studied kinases (light kinases), while the training primarily comprises under-studied kinases (dark kinases). However, it’s crucial to note that this arrangement is solely for evaluation purposes; the deployed model can predict dark kinases.
ii. **Stratification based on kinase groups:** Kinases within the same kinase group share evolutionary relationships and functional similarities [Manning et al., 2002]. After preprocessing, the dataset contains only 392 kinases distributed across 11 kinase groups and 129 kinase families. Stratifying by kinase families is impractical due to the limited number of kinases per family, which would hinder equal representation of each kinase group in each split. Thus, we stratify kinases based on their group membership, ensuring the representation of kinase groups in the training, validation, and test sets whenever possible.
iii. **Sequence similarity of kinases:** In the light of the inference task, which is to predict the kinase for a given phosphosite, to avoid optimistic performance estimates, kinases with sequence similarity are grouped and assigned exclusively to the same set (train, validation, or test). This criterion is important to prevent the model from being trained on kinases that are highly similar to the kinases in the test set, thereby avoiding optimistic evaluation results. It also aligns with the principles of zero-shot learning by guaranteeing that all kinases in the test set are entirely new to the model. Sequence similarity is determined by sequence identity calculated after pairwise global alignment of the kinase domains.

Note that a single phosphosite can be targeted by multiple kinases, which may result in the same phosphosite appearing in both the training and test sets with different kinase labels. We quantified the multi-label nature of the task in Supplementary Figure S3, which shows the number of sites associated with a single kinase or multiple kinases in each split. Additionally, we report the sites unique to the test set or shared with the validation and training data in the Supplementary Figure S4. As the aim is to predict the right kinases for a known phosphosite, a phosphosite-based split of the train, validation, and test set is not necessary. Even when a phosphosite appears in both splits, the associated kinase labels are disjoint across training and test sets. The model is still required to generalize to unseen kinases. It is indeed more challenging for the model, as it has previously associated this site with a different kinase and now needs to predict its association with the unseen test kinase. Therefore, not splitting based on the phopshosites does not affect the integrity of the evaluation process.

Taking all these aspects into consideration, we divided the dataset into training (80%), validation (10%), and test (10%) sets. We first categorize kinases as train or test kinases according to the number of phosphosites they are associated with. Kinases that are associated with fewer than 15 phosphosites are defined as train kinases. Later, kinases with at least 90% sequence identity similarity are grouped and are randomly defined as entirely train or test kinases altogether. From the remaining kinases, test kinases are randomly selected from each kinase group in a stratified manner to ensure sufficient test example pairs from each kinase group. All remaining kinases are designated as train kinases. This process is repeated to determine validation kinases from among train kinases by setting the threshold for kinases in validation to be at least 10 phosphosites per kinase. Finally, the train, validation, and test sets include all train phosphosite-kinase pairs associated with the kinases in that relative set. Splitting the kinases into train, validation, and test is performed in a randomized and reproducible manner. Thus, different splits of the DARKIN dataset can be generated by setting different random seeds.

We evaluate our methods using the macro Average Precision (AP) score. AP summarizes the precision-recall curve at all recall levels[Salton and McGill, 1983]. In this way, AP provides a measure of how well the model is able to rank positive samples over negative samples. By using AP, for each kinase, we rank the prediction probabilities made for all phosphosite samples. If the model is able to assign relatively higher probabilities to phosphosites that are actually known to be phosphorylated by that kinase (the ground-truth phosphosites for that kinase), then we obtain higher scores closer to 1, indicating that the model ranks positive sites above negative ones, and hence achieves higher AP scores. In our setup, we calculate the AP score for each kinase and then take the mean across all kinase classes, hence calculating the macro AP. Although top-k accuracy is a well-known metric, in our setting it fluctuated sharply – small changes in predictions for the sparsely represented kinase classes produced large jumps in the score. To counter this instability and the effects of class imbalance, we report *macro* average precision, which assigns equal weight to each kinase class. macro AP, therefore, provides a steadier assessment of performance across both common and rare classes. When multiple kinases can phosphorylate a phosphosite, we accept the predicted kinase as a true positive if it matches any of the true kinases associated with it.

### Zero-Shot Classifiers

We employ two zero-shot learning (ZSL) models in our experiments. The first is a fitting-free method based on an adapted k-NN classifier, intentionally kept simple. The second model is a well-established bilinear zero-shot compatibility model. Further details on these approaches are provided in the following sections.

#### Zero-Shot k-NN Classifier

To benchmark the zero-shot dark kinase prediction performance, we devised a simple baseline method by adapting the principles of the k-NN algorithm for supervised classification to our zero-shot classification task. For a given test phosphosite, we first locate the *k* most similar training phosphosites in the phosphosite representation space. Subsequently, we identify the most common *light kinase* among the kinases associated with the nearest neighbor phosphosites. In cases where there is no majority, we choose the nearest neighbor’s light kinase. Unlike the supervised k-NN approach, we predict the dark kinase that resembles the most to this predicted light kinase in the representation space. Kinase similarity is assessed using the cosine similarity of the kinase embedding vectors. These cosine similarity scores are considered our prediction scores, indicating how likely each dark kinase is to phosphorylate the test phosphosite at hand. This procedure is depicted in Figure 1a.

**Fig. 1.**
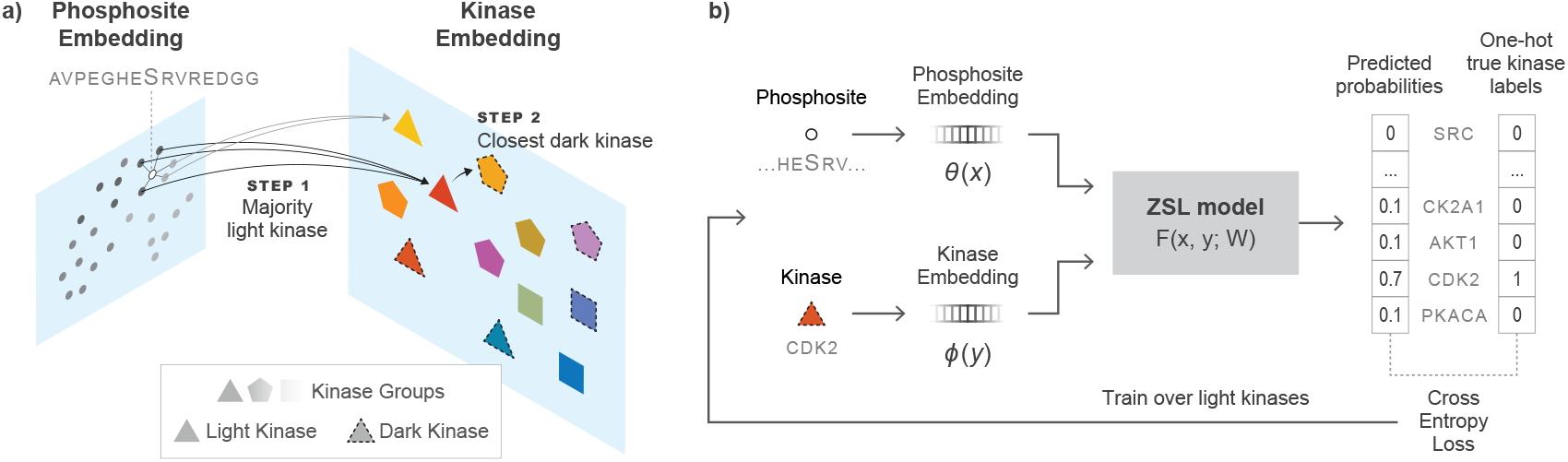
a) k-NN based zero-shot classifier. First, the test phosphosite’s nearest neighbor phosphosites are determined in the training data. The majority vote is taken among the neighbors’ class labels to pick the most likely light kinase. Then, the dark kinase most similar to this light kinase is picked. b) The bilinear compatibility function *F* takes the phosphosite and kinase embedding vectors and is trained to minimize the cross-entropy loss over light kinases. At the prediction time, *F* is used to assess the compatibility of the phosphosite and the dark kinases.

Our motivation for devising this method is to evaluate the pLMs as *directly* as possible, in the sense that the approach does not involve numerical optimization, and the only hyperparameter is *k*. This simplicity provides an additional view of the relative strengths of the pLMs, largely avoiding model selection effects.

#### Bilinear Zero-Shot Learning Model

The second zero-shot learning method we use is a bilinear compatibility model. While a variety of other zero-shot learning methods, particularly in image classification, have been proposed over the years, variants based on bilinear compatibility models are arguably among the most established [Xian et al., 2017, Akata et al., 2016, Romera-Paredes and Torr, 2015, Frome et al., 2013, Akata et al., 2015, Kodirov et al., 2017, Sumbul et al., 2018, Deznabi et al., 2020]. Therefore, they are particularly suitable for our pLM evaluation purposes.

The bilinear zero-shot model (BZSM) aims to estimate the compatibility between a given pair of phosphosite *x* and kinase *y* (illustrated in Figure 1b). In our work, we use the formulation variant proposed and used in [Sumbul et al., 2018, Deznabi et al., 2020], which defines the compatibility function *F* (*x, y*) = [*θ*(*x*)^⊤^ 1]*W* [*ϕ*(*y*)^⊤^ 1]^⊤^ where *θ*(*x*) ∈ ℝ^*d*^ is the phosphosite representation, and *ϕ*(*y*) ∈ ℝ^*m*^ is the kinase representation.

The augmentation of both representations with separate bias dimensions increases the expressivity of the model [Sumbul et al., 2018], which can more clearly be observed when the definition is expanded:

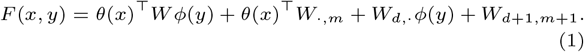

In this formulation, the first term estimates pairwise compatibility. The second term acts analogous to a log *p*(*x*) prior, formulated via a linear estimator conditioned on *θ*(*x*). Similarly, the third term is a log *p*(*y*) prior, expressed as a linear function of *ϕ*(*y*). And finally, the last term is simply a trainable scalar. The model is trained by minimizing the regularized cross-entropy loss:

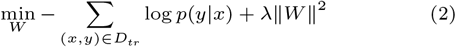

where the summation runs over all phosphosite-kinase pairs available in the training set *D*_*tr*_ = (*x*_*i*_, *y*_*i*_), and *p*(*y*|*x*) is the softmax of *F* over the light kinases:

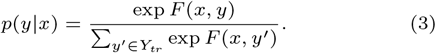

The *ℓ*_2_ regularization term in Eq. 2 is implemented as *weight decay* in practice. At test time, *p*(*y*|*x*) is calculated via softmax over the test kinases.

## Results

### Hyperparameter tuning

We use macro AP on the validation set for model selection in all cases. For k-NN based ZSL, we choose *k* from {3, 5, 7}. For the bilinear ZSL, we perform a hyperparameter search among random combinations of learning rate (0.000001…0.1), optimizer (Adam, SGD, RMSprop), learning rate schedule (Exponential, Step, CosineAnnealing), momentum (0.95…0.9999) and the weight decay (0.00001…0.01). Finally, to measure the effect of initialization, unless otherwise stated, we train BZSM models three times and report the mean and standard deviation of the macro AP values.

### DARKIN benchmark statistics

We present four DARKIN splits^∗^ for researchers. The experiments are conducted using DARKIN Split 1 unless otherwise specified. Therefore, we share the statistics for Split 1. The number of kinases, distinct phosphosites, and the phosphosite-kinase associations in the train, validation, and test sets are shown in Figure 2. Furthermore, the histogram displaying the number of kinases associated with specific numbers of phosphosites is presented in Figure 3. The balanced distribution of kinases according to kinase groups and the resulting kinase-phosphosite pair distribution can be analyzed in Supplementary Figure S2, which results from the stratification strategy we used when splitting the kinase-phosphosite pairs. In addition to these statistics, further statistics such as the number of single-kinase and multi-kinase phosphosites (Supplementary Figure S3), seen and novel sites in the test dataset (Supplementary Figure S4), and the distribution of sites by the number of kinases they are associated with in the train, validation, and test sets (Supplementary Figure S5) are accessible in the Supplementary Text.

**Fig. 2.**
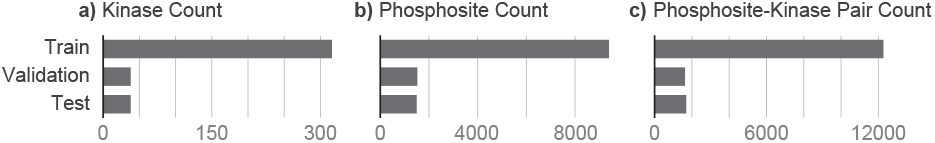
a) The number of kinases b) The number of unique phosphosites c) The number of kinase-phosphosite pairs in each train, validation, and test folds of the default DARKIN split dataset.

**Fig. 3.**
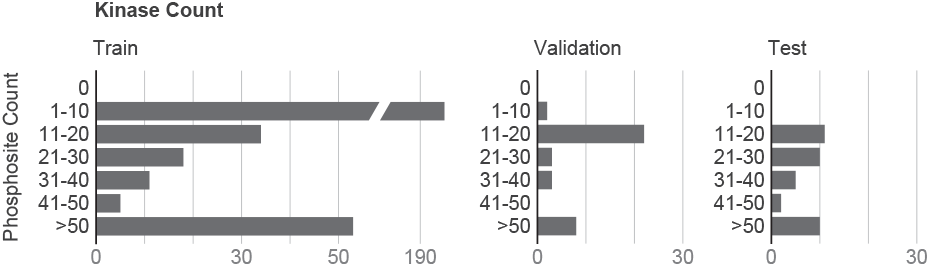
The histogram of the number of phosphosites associated with kinases in train, validation, and test sets in the default DARKIN split. See Section 2.4 for details.

### Comparison of Protein Language Models

We initially assess the effectiveness of pLM-based embeddings using both k-NN and BZSM methods. Table 2 presents macro AP scores obtained through the k-NN and BZSM when different pLM embeddings (detailed in Table 1) are used to represent the 15-mer around the phosphosite sequence and the kinase domain sequence. When employing pLM embeddings, we utilize embeddings sourced from the same pLM for both the phosphosite and kinase. To establish baseline performance, we also present results obtained with three sequence encoding methods: one-hot encoding, BLOSUM62, and NLF encoding (Section 2.3). In both models, we observe that most pLM representations outperform the baseline encodings, indicating that they capture the protein sequences’ relevant characteristics better.

**Table 2.**
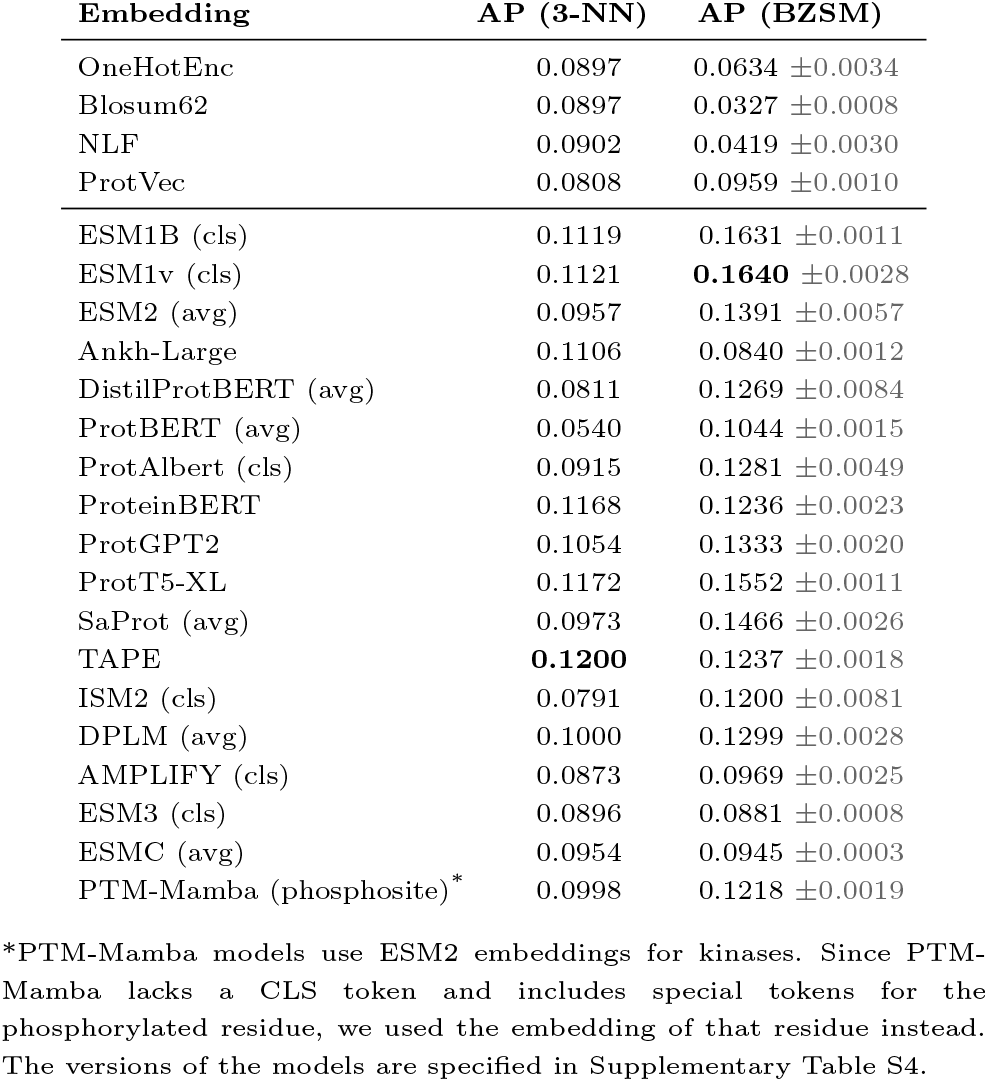
Mean macro AP of 3-NN and Bilinear Zero-Shot Model (BZSM) using only pLM embeddings. For pLM with CLS versus average token embedding alternatives, the best performing one is shown.

The TAPE embeddings perform the best among the k-NN models (0.12 AP score). The ESM models and ProtT5-XL closely follow TAPE’s results (Table 2). In the BZSM models, however, the TAPE embeddings fall behind the ESM1B and ESM1v embeddings. The superior performance of TAPE in the k-NN could be due to it being a lower-dimensional vector (see Table 1). In BZSM, when employing the CLS token, ESM1B and ESM1v achieve over 0.16 macro AP. ProtT5-XL is the third close runner-up, and SaProt (cls) also performs well.

### CLS token embedding versus averaging

Several pLMs provide a CLS token whose embedding is commonly used as the sequence summary [Devlin et al., 2019]. However, it is not clear whether the CLS token or the average of all token embeddings provides a better summary for this task. The performance differences between these two alternatives are shown in Figure 4, indicating that (i) the results can depend on this detail, and (ii) the right option varies across the pLMs.

**Fig. 4.**
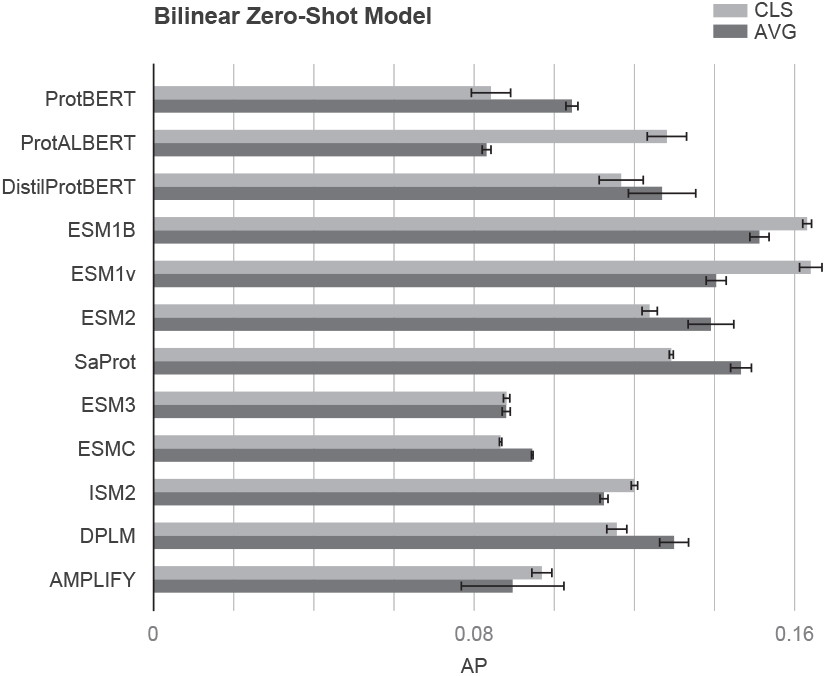
Performance comparison of BZSM models across different pLMs. The x-axis represents the Average Precision (AP) and the y-axis lists the evaluated pLMs. Light gray bars correspond to results obtained using the CLS token representation, while dark gray bars correspond to results obtained using the average of all token embeddings. Error bars indicate standard deviation across multiple runs.

### Incorporating additional kinase information

We augment the kinase sequence embedding vectors with additional information regarding kinase family hierarchy and EC classification. We encode these memberships as one-hot encoded vectors and append them to the sequence embedding vectors. Here, we experiment only with the BZSM since it outperforms the k-NN (The complete results obtained on the 3-NN with this additional kinase information are provided in Supplementary Table S1). Including each type of additional information individually enhances the performance of all models (Table 3), especially the inclusion of the kinase family information. The models based on ESM1B, ESM1v, and SaProt, using the CLS token embeddings, benefit the most and emerge as the best performers in this augmented case. These findings underscore that there is additional information in these kinase categorizations that cannot be captured solely with sequence information. The detailed list of results obtained with all pLMs obtained on the BZSM with this additional kinase information is provided in Supplementary Table S2.

**Table 3.** The BZSM performance trained with sequence embedding and other kinase information. The mean macro APs are shown. Of CLS and embedding averaging, only the best-performing model results are listed.

| Embedding | Base | + Family | + Group | + EC | + Family + Group + EC |
| --- | --- | --- | --- | --- | --- |
| OneHotEnc | 0.0634 | 0.1107 | 0.0832 | 0.0802 | 0.1098 |
| Blosum62 | 0.0327 | 0.0318 | 0.0310 | 0.0337 | 0.0323 |
| NLF | 0.0419 | 0.0391 | 0.0425 | 0.0400 | 0.0426 |
| ProtVec | 0.0959 | 0.1262 | 0.1129 | 0.1214 | 0.1354 |
| ProtBERT (cls) | 0.0842 | 0.1170 | 0.1077 | 0.1132 | 0.1273 |
| ProteinBERT | 0.1236 | 0.1506 | 0.1215 | 0.1367 | 0.1359 |
| ProtT5-XL | 0.1552 | 0.1701 | 0.1531 | 0.1674 | 0.1731 |
| ESM1B (cls) | 0.1631 | **0.1740** | **0.1688** | **0.1680** | 0.1769 |
| ESM1v (cls) | **0.1640** | 0.1737 | 0.1653 | 0.1652 | 0.1734 |
| ESM2 (avg) | 0.1391 | 0.1588 | 0.1453 | 0.1496 | 0.1638 |
| DistilProtBERT (cls) | 0.1167 | 0.1360 | 0.1292 | 0.1287 | 0.1441 |
| ProtGPT2 | 0.1333 | 0.1476 | 0.1412 | 0.1419 | 0.1557 |
| Ankh-Large | 0.0840 | 0.1417 | 0.1135 | 0.1178 | 0.1594 |
| ProtAlbert (cls) | 0.1281 | 0.1269 | 0.1276 | 0.1285 | 0.1372 |
| SaProt (cls) | 0.1292 | 0.1696 | 0.1424 | 0.1434 | **0.1800** |
| TAPE | 0.1237 | 0.1379 | 0.1333 | 0.1310 | 0.1455 |
| ISM2 (cls) | 0.1200 | 0.1275 | 0.1260 | 0.1333 | 0.1374 |
| DPLM (avg) | 0.1299 | 0.1427 | 0.1318 | 0.1368 | 0.1420 |
| AMPLIFY (avg) | 0.0896 | 0.0968 | 0.0944 | 0.0969 | 0.1066 |
| ESM3 (cls) | 0.0881 | 0.1484 | 0.1220 | 0.1238 | 0.1611 |
| ESMC (cls) | 0.0866 | 0.1672 | 0.1136 | 0.1401 | 0.1754 |
| PTM-Mamba (phosphosite)* | 0.1218 | 0.1432 | 0.1292 | 0.1346 | 0.1471 |

\*PTM-Mamba models utilize ESM2 embeddings for kinases. Since PTM-Mamba lacks a CLS token and includes special tokens for the phosphorylated residue, we used the embedding of that residue instead.

### Comparing the best-performing pLMs on different random DARKIN splits

As ESM1B and SaProt emerge as the two top-performing pLMs when paired with the BZSM model (Table 3), we further evaluated their performance on three additional random splits of the DARKIN dataset to facilitate a more comprehensive comparison between these two pLMs. While both models demonstrate competitiveness, SaProt consistently outperforms ESM1B slightly across all runs on these four different splits (Table 4). The performance of SaProt underscores the added value of structural information.

**Table 4.** The mean macro AP scores at multiple levels (family, group, phosphosite) for the two best-pLMs, ESM1B (Family + Group + EC) and SaProt (Family + Group + EC), on four random DARKIN splits for the BZSM.

| Split | Embedding | AP | Phospho-site AP | Family AP | Group AP | Masked Group AP |
| --- | --- | --- | --- | --- | --- | --- |
| Split 1 | ESM1B (cls) | 0.1769 | 0.2830 | 0.2278 | **0.3959** | **0.4054** |
|  | SaProt (cls) | **0.1800** | **0.3053** | **0.2384** | 0.3903 | 0.3868 |
| Split 2 | ESM1B (cls) | 0.1536 | 0.2747 | 0.1989 | **0.3689** | 0.3644 |
|  | SaProt (cls) | **0.1599** | **0.2929** | **0.2087** | 0.3649 | **0.3702** |
| Split 3 | ESM1B (cls) | 0.1531 | 0.2951 | 0.1987 | **0.3747** | 0.3508 |
|  | SaProt (cls) | **0.1627** | **0.3142** | **0.2104** | 0.3663 | **0.3598** |
| Split 4 | ESM1B (cls) | 0.1652 | 0.3118 | 0.2142 | 0.3969 | 0.3563 |
|  | SaProt (cls) | **0.1690** | **0.3482** | **0.2205** | **0.4069** | **0.3674** |

### Extended evaluation of kinase family, group, and phosphosite predictions

We evaluated model performance on all DARKIN splits using macro AP at multiple levels: Family AP, where kinase predictions are aggregated by their families; Group AP, where kinase predictions are aggregated by their groups; and Phosphosite AP, which evaluates precision for phosphosite-specific predictions. We also calculated the hit@k accuracy for these models (Supplementary Table S3). In all these metrics, SAProt shows slightly better performance.

Additionally, we introduced a metric, Masked Group AP, which focuses on precision within the true group by excluding irrelevant kinases from the predictions. This metric works by masking logits for kinases outside the ground truth kinase group, effectively setting them to negative infinity. This metric simulates a scenario where the model perfectly identifies kinase groups and predicts within the group. This allows us to measure the model’s ability to rank kinases accurately within groups. Our findings, summarized in (Table 4), show that Masked Group AP significantly outperforms standard AP, with values greater than twice those of standard AP calculated over all classes. This improvement demonstrates the strong impact of incorporating group-level information, suggesting that if kinase groups could be predicted accurately—whether by this or a separate model—the performance jump in kinase ranking could be substantial. This insight suggests a promising direction for future research, where accurate group predictions could serve as a basis for refining kinase-level predictions.

### Fine-tuning of Phosphosite and Kinase Encoders

To evaluate if task-specific fine-tuning improves the performance, we extended the original BZSM setup—which keeps phosphosite and kinase embeddings fixed and only learns the compatibility matrix—by adding four fine-tuning variants and evaluating them using the two well-performing pLMs, ESM1B and ProtT5-XL. First, we allowed end-to-end fine-tuning of the phosphosite encoder while keeping the kinase encoder frozen and still learning the compatibility matrix W. Next, we gradually unfroze the kinase encoder, reinitializing and training either its final transformer block or, in a deeper variant, the last two blocks, so that both phosphosite and kinase representations could adapt jointly with the compatibility matrix *W* . Finally, in the fourth variant we experimented with a fully shared encoder that produces both phosphosite and kinase embeddings; here, the entire model is fine-tuned jointly, and compatibility is computed via a simple dot product, eliminating the need for W. Each regime was tested with two kinase representations: sequence-only embeddings and appending the sequence embeddings with family, group, and EC information vectors.

As presented in Table 5, fine-tuning the pLM encoders does not guarantee improved performance. Instead, the results were inconsistent across different configurations. For the ESM1b model, the highest performance was AP of 0.1911, achieved by reinitializing the final two layers of both the phosphosite and kinase encoders using the full set of kinase features. However, this represents only a marginal improvement over other configurations and comes at a notable computational cost. Similarly, the ProtT5-XL model saw a slight performance increase to an AP of 0.1800 when reinitializing the last layer of both encoders. Notably, most other fine-tuning strategies resulted in a decrease in performance for both models.

**Table 5.**
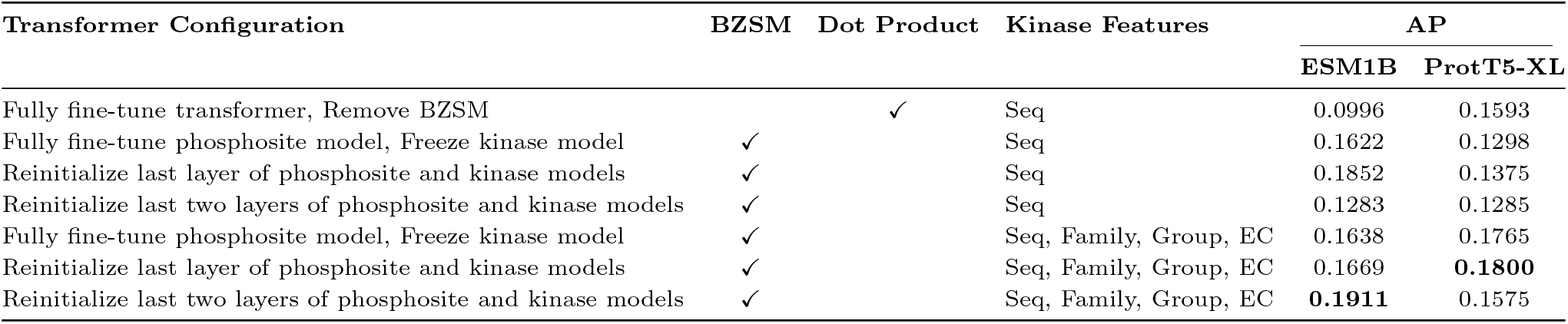
Experiments on fine-tuning ESM1b and ProtT5-XL, in which we employ transformers to fine-tune either the phosphosite model or the phosphosite and kinase model simultaneously. As a baseline evaluation, we remove BZSM and evaluate zero-shot predictions by the dot product of learned kinase and phosphosite representation. ESM1b embeddings are obtained using CLS token representation, while ProtT5-XL embeddings are obtained using the average of all token embeddings. Training and evaluation protocols are the same for both pLMs.

We explored other fine-tuning strategies. To arrive at phosphosite-aware and kinase-aware pLMs, we conducted comprehensive experiments in which we fine-tuned PLMs with kinase- and phosphorylation-related auxiliary tasks. These tasks include i) phosphorylation prediction, given a potential phosphosite and its surrounding sequence, the model is trained to predict if this site is phosphorylated or not. ii) Kinase group prediction, given the kinase domain sequence, predicting the group of the given kinase. This is a multi-class classification task. iii)Contrastive learning on family/group relations. In this task, the model should learn the kinase family/group relationships in a contrastive learning setup. We present the dataset, experimental methods, and the results in the Supplementary Text Section 4. None of these phosphosite and kinase fine-tuning strategies match the performance of the end-to-end fine-tuning presented above (AP score of 0.1911) obtained by reinitializing the last two layers of the ESM1B model.

## Conclusion

Focused on the zero-shot task of assigning phosphosites to understudied dark kinases, DARKIN offers a novel benchmark for evaluating pLMs. As it is easy to fall into the data leakage pitfalls in these types of problems, as raised and discussed in drug-target prediction [Chatterjee et al., 2023], drug synergy prediction [Çandır et al., 2025], in genomics [Whalen et al., 2022], or link prediction [Brière et al., 2025], it is important to evaluate the models in robust evaluation frameworks to assess the generalization of these models [Bernett et al., 2024]. In this work, the train, validation, and test splits are carefully designed to follow zero-shot learning and kinase-related issues. We evaluate the pLMs’ representation capabilities in this problem using two zero-shot classifiers. Our results demonstrate the superior performance of the ESM models, the ProtT5-XL, and the SaProt models.

Based on our results using the DARKIN dataset, dark kinase–phosphosite prediction remains a highly challenging task for the current pLMs. The highest AP score achieved was 0.1911 using fine-tuning pLMs, which considerably outperforms random guessing (0.03 by averaging AP over 1000 runs of randomly generated ranking of kinases for a given site), but can be considered low overall. The low performance could be due to several reasons. There are challenging cases where the phosphosite sequences are almost identical, but the associated kinase sets for these phosphosites differ. This difference could be due to a true biological difference that can be explained by a structural or functional difference (a required interaction partner or the same cellular localization), or it could also be an issue of data incompleteness. While some kinase-phosphosite pairs are truly associated, they might not have been experimentally studied and therefore are not reported as associated pairs. We should also note that the performances in a deployed model of dark-kinase associations are likely to be higher. To ensure a sufficient number of examples in the evaluation, as explained in the Method section, we switched the light and dark kinases in the train and test sets. In this way, the test set included the well-studied kinases with more examples, and the training set included the understudied kinases. While this strategy is useful for benchmarking purposes, it poses a challenge in training, as the training data contains many kinases with few examples. Since the deployed model uses the well-studied kinases as well, it is likely to have better predictive performance.

In this study, we excluded fusion kinases and non-canonical kinase isoforms in constructing the datasets. This was due to the lack of annotation of their kinase domains in some cases and the low number of known associated phosphosites, which made it difficult to reliably evaluate the models’ performance on these kinases. These kinase forms can play crucial roles in disease contexts such as cancer, where gene fusions or isoform specific events give rise to novel or dysregulated signaling activities [Stransky et al., 2014, Gonzalez and McGraw, 2009]. Thus, zero-shot predictions coupled with experimental validation on these kinases can open new avenues for understanding the functional impact of isoforms and oncogenic fusions.

The study focused on the zero-shot learning framework. Another promising direction and interesting benchmark is the few-shot learning problem, in which the model leverages the few known phosphosites of the kinases during the training. The current DARKIN dataset can be modified for this setup easily. We hope this novel benchmark will facilitate comprehensive evaluations of pLMs and dark kinase prediction models, contributing to protein biology research.

## Acknowledgments

This work was supported by the Scientific and Technological Research Council of Turkey (TUBITAK) under Grant #122E500. The numerical calculations reported in this paper were fully/partially performed at TUBITAK ULAKBIM, High Performance and Grid Computing Center (TRUBA resources).

## Footnotes

∗ https://github.com/tastanlab/darkin https://zenodo.org/records/16729884

