## Supplementary Information for "DARKIN: A zero-shot benchmark for phosphosite–dark kinase association using protein language models"

### Supplementary Materials for "DARKIN: A zero-shot classification benchmark and an evaluation of protein language models"

Emine Ayşe Sunar[1,†], Zeynep Işık[1,†], Mert Pekey[1,†], Ramazan Gokberk Cinbis[2,*], and Oznur Tastan[1,*]

[1]Sabanci University, Department of Computer Science and Engineering, İstanbul, Türkiye
[2]Middle East Technical University, Department of Computer Engineering, Ankara, Türkiye
[†]These authors contributed equally to this work.

#### 1 Darkin Dataset Splitting Algorithm

To construct the DARKIN datasets, we implemented a reproducible and configurable dataset-splitting algorithm for the zero-shot learning setup. The important parameters of the algorithm, along with their description, are listed in Table S1, and the full list is available in the GitHub repository: `https://github.com/tastanlab/darkin`.

The steps of the algorithm are as follows (assuming both `TAKE_SEQUENCE_SIMILARITY_INTO_CONSIDERATION` and `DIVIDE_WRT_GROUP` are set to `True`):

1. First, for each kinase, the number of unique kinase–phosphosite associations is calculated.
2. The total number of kinase–phosphosite associations per kinase group is computed by summing the counts from the previous step across all kinases belonging to the same group.
3. Based on the parameter `STRATIFY_PERCENTAGE_FOR_UNSEEN_TEST_KINASE`, the number of phosphosite–kinase associations that should be allocated to the test set is computed per group. Note that this step only determines the counts to be added to the test set, not the specific kinase-phosphosite associations.
4. Next, kinases are grouped based on sequence similarity, and these clusters are used to guide the assignment of kinases to the train and test sets:

| Parameter | Description |
|---|---|
| RANDOM_SEED | Random seed for reproducibility. Used to generate deterministic splits. |
| KINASE_SIMILARITY_ PERCENT | Threshold for kinase domain sequence identity. Kinases with similarity above this value are grouped into the same split. |
| KINASE_COUNT_ TEST_THRESHOLD | Minimum number of phosphosite associations required for a kinase to be included in the test set. |
| STRATIFY_PERCENTAGE_ FOR_UNSEEN_ TEST_KINASE | The percentage of the dataset that should be entered into the test set as unseen data. |
| INCLUDE_VALIDATION | Boolean flag indicating whether to generate a validation set from the training split. |
| TAKE_SEQUENCE_ SIMILARITY_INTO_ CONSIDERATION | If `True`, groups similar kinases in the same split to prevent information leakage. |
| DIVIDE_WRT_GROUP | Whether to stratify kinases by group label when splitting. If set to `False`, kinase group distributions may be imbalanced. |

Table S1: A subset of the parameters used in the dataset splitting algorithm. The full list of parameters can be found in our GitHub repository: `https://github.com/tastanlab/darkin`

- Before assigning kinases to any split, kinase clusters are constructed such that each cluster contains kinases whose domain similarity is equal to or greater than the threshold set by `KINASE_SIMILARITY_PERCENT`.
- Each cluster is processed independently. If any kinase in a cluster has fewer associations than `KINASE_COUNT_TEST_THRESHOLD`, all of the kinases in that cluster is assigned to the training set.
- If none of the kinases in the cluster have fewer phosphosite associations than the threshold, then with probability $\theta$ the cluster is assigned to the test set, and with probability $1-\theta$ to the training set. We set $\theta = 0.6$ empirically with the aim of achieving a balanced distribution between the training and test sets.

5. After processing the similarity clusters, any remaining kinase that has fewer phosphosite associations than `KINASE_COUNT_TEST_THRESHOLD` is assigned to the training set.
6. For the rest of the kinases (i.e., those eligible for either split and not part of already-assigned similarity clusters), kinases are added to the test set until the precomputed number of kinase–site associations (from Step 3) per group is fulfilled. Once this target is met, all remaining kinases are added to the training set.
7. At this point, the number of sites that should be assigned to the train and tests is known. Next, the kinase–phosphosite association rows are distributed between the train and test sets

based on the number of samples determined for each kinase. Since a phosphosite may be associated with multiple kinases, the following rules are applied for each kinase-phosphosite pair:

- If a phosphosite is only associated with training kinases, it is added to the training set.
- If a phosphosite is only associated with test kinases, it is added to the test set.
- If a phosphosite is associated with both training and test kinases:
  - A version of the sample containing only the training kinases is added to the training set. (e.g., if a site is associated with both training and test kinases, the training set will include that site paired only with the training kinases)
  - A version of the same sample containing only the test kinases is added to the test set (i.e., the site is duplicated and paired only with its test kinases in the test set).

8. The above procedure is applied until the required number of kinase–phosphosite association samples is reached for each kinase group in the test set. All remaining samples are added to the training set.

9. The same logic used in Steps 1–8 is then applied to the training set to generate a validation set if the parameter `INCLUDE_VALIDATION` is set to `True`. After this step, the complete `train-validation-test` split is finalized.

We explain this procedure in Algorithm 1. Let the input be the full set of kinase–phosphosite associations and the following configurable parameters: `RANDOM_SEED`, `KINASE_SIMILARITY_PERCENT`, `KINASE_COUNT_TEST_THRESHOLD`, `STRATIFY_PERCENTAGE_FOR_UNSEEN_TEST_KINASE`, `INCLUDE_VALIDATION`, `TAKE_SEQUENCE_SIMILARITY_INTO_CONSIDERATION`, and `DIVIDE_WRT_GROUP`.

**Algorithm 1** DARKIN zero-shot dataset splitting

**Require:** Kinase–phosphosite association data, parameters. $\theta$: test-set assignment probability for kinase similarity clusters.

**Ensure:** Train/(optional) validation/test splits with zero-shot constraints, group stratification, similarity constraints.

Set random seed to `RANDOM_SEED`.<br>
For each kinase $k$: $A_k \leftarrow$ number of unique phosphosite associations.<br>
**if** `DIVIDE_WRT_GROUP` **then**<br>
  For each group $g$, compute $T_g \leftarrow \sum_{k \in g} A_k$.<br>
  $S_g \leftarrow \lfloor T_g \times$ `STRATIFY_PERCENTAGE_FOR_UNSEEN_TEST_KINASE`$\rfloor$.<br>
**else**<br>
  Treat full dataset as single group with analogous $T, S$.<br>
**end if**<br>
**if** `TAKE_SEQUENCE_SIMILARITY_INTO_CONSIDERATION` **then**<br>
  Build clusters $\mathcal{C}$: kinases with pairwise identity $\geq$ `KINASE_SIMILARITY_PERCENT`.<br>
**else**<br>
  Each kinase is its own cluster.<br>
**end if**<br>
Initialize $\mathcal{K}_{\text{train}}, \mathcal{K}_{\text{test}} \leftarrow \emptyset$.<br>
**for all** cluster $C \in \mathcal{C}$ **do**<br>
  **if** $\exists k \in C$ with $A_k <$ `KINASE_COUNT_TEST_THRESHOLD` **then**<br>
    $\mathcal{K}_{\text{train}} \mathrel{+}= C$<br>
  **else**<br>
    Sample $u \sim \text{Uniform}(0, 1)$.<br>
    **if** $u < \theta$ **then**<br>
      $\mathcal{K}_{\text{test}} \mathrel{+}= C$<br>
    **else**<br>
      $\mathcal{K}_{\text{train}} \mathrel{+}= C$<br>
    **end if**<br>
  **end if**<br>
**end for**<br>
Assign any remaining kinase $k$ with $A_k <$ `KINASE_COUNT_TEST_THRESHOLD` to $\mathcal{K}_{\text{train}}$.<br>
**Group-aware stratification:** For each group $g$, while test associations from $\mathcal{K}_{\text{test}} \cap g < S_g$, randomly select unassigned $k \in g$ with $A_k \geq$ `KINASE_COUNT_TEST_THRESHOLD` and move it to $\mathcal{K}_{\text{test}}$.<br>
Assign all remaining unassigned kinases to $\mathcal{K}_{\text{train}}$.<br>
**Distribute kinase–phosphosite samples:**<br>
**for all** phosphosite $s$ **do**<br>
  Let $K_{\text{train}}(s)$ / $K_{\text{test}}(s)$ be associated kinases.<br>
  **if** $K_{\text{train}}(s) \neq \emptyset$ and $K_{\text{test}}(s) = \emptyset$ **then**<br>
    Add $(s, K_{\text{train}}(s))$ to training.<br>
  **else if** $K_{\text{test}}(s) \neq \emptyset$ and $K_{\text{train}}(s) = \emptyset$ **then**<br>
    Add $(s, K_{\text{test}}(s))$ to test.<br>
  **else**<br>
    Add both: training $(s, K_{\text{train}}(s))$ and test $(s, K_{\text{test}}(s))$.<br>
  **end if**<br>
**end for**<br>
**Finalize:** Ensure test set meets $S_g$; leftovers remain in train.<br>
**if** `INCLUDE_VALIDATION` **then**<br>
  Recursively apply above to split the validation set.<br>
**end if**<br>
**return** splits.

#### 2 Dataset Statistics

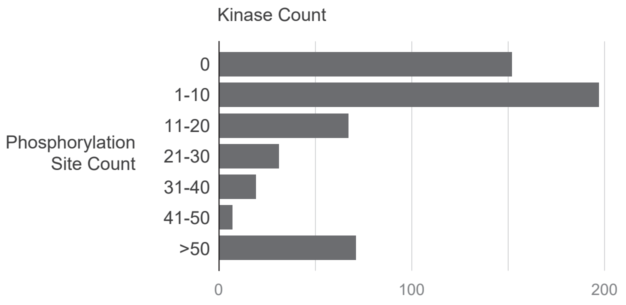


Figure S1: The histogram of the number of phosphosites associated with a kinase in the PhosphositePlus dataset. For most of the kinases, no or few phosphosites are assigned.

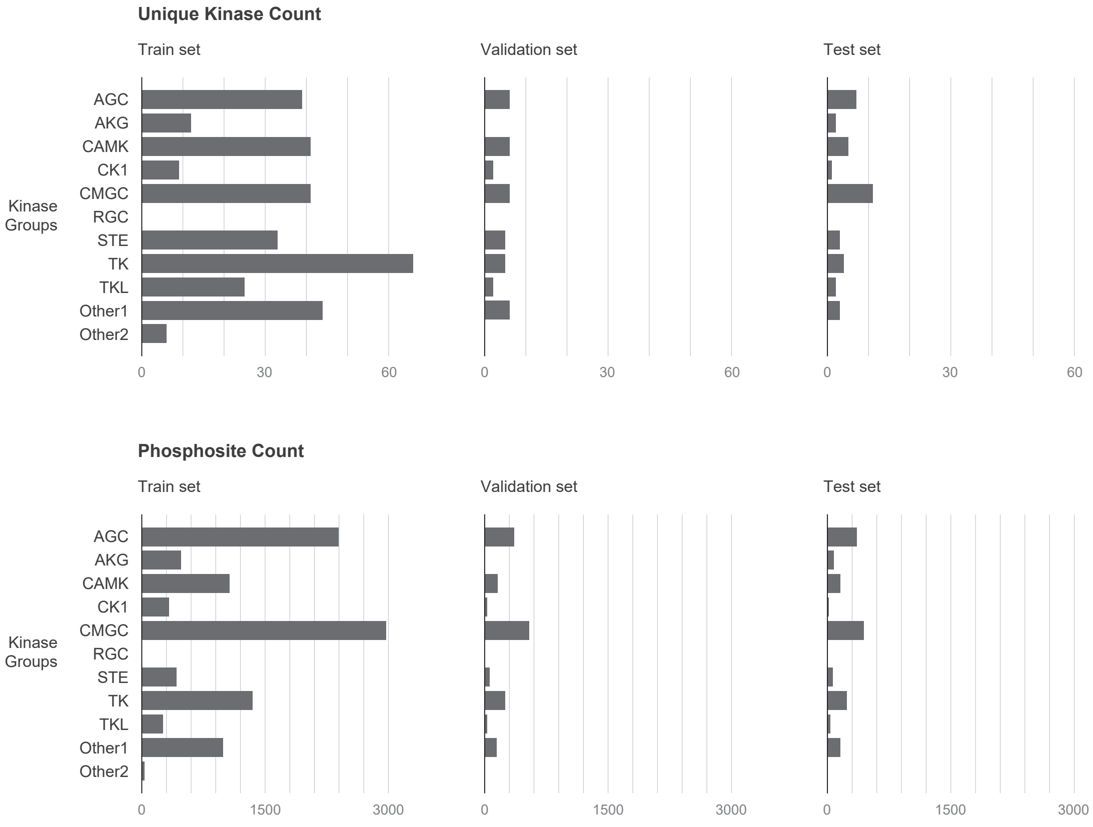


Figure S2: Splits are stratified based on kinase groups. (Upper panel) The number of kinases in each kinase group, in train, validation, and test folds. (Lower panel) The number of unique phosphosites that are associated with a kinase that falls into the kinase group in train, validation, and test folds.

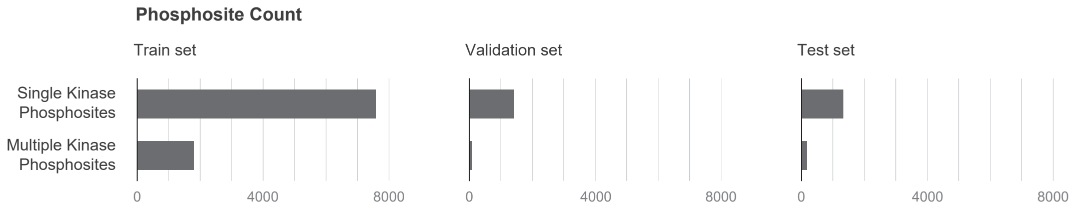


Figure S3: The number of multi-label cases for train, validation, and test. In training, we binarize the multi-labels; in testing, we count the prediction as a true positive if the predicted kinase is in the set. The number of multi-label cases is small in the test set.

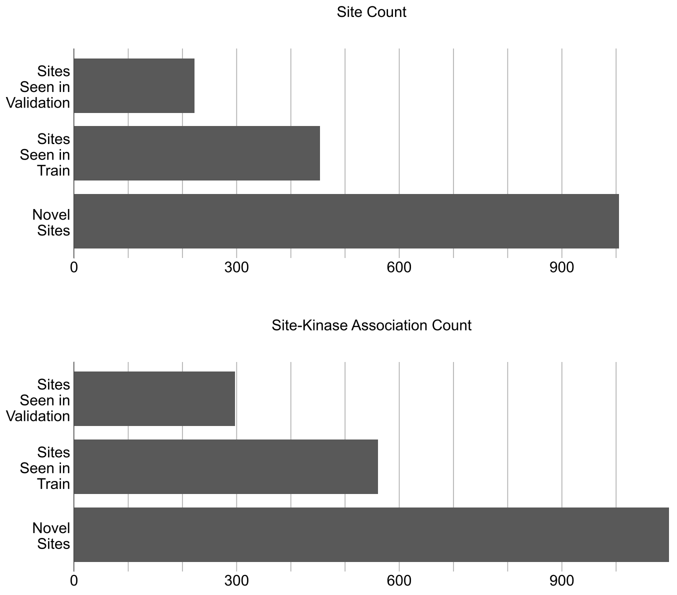


Figure S4: (Top figure): The count of novel sites in the test split, and the count of the common sites of the test split with the train and validation splits. (Bottom Figure): The count of the kinase-site association counts related to the novel sites in the test split, and likewise the count of the kinase-site association counts related to the common sites of the test split with the train and validation splits.

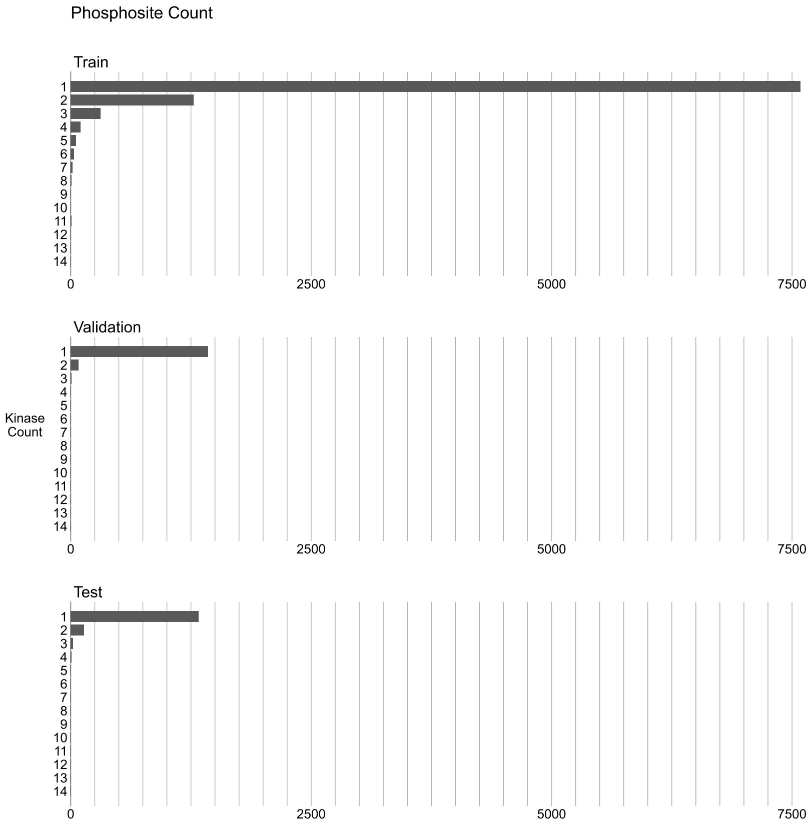


Figure S5: The histogram of the number of kinases associated with the sites. This plot could be read like this: “In the train split, there is around 1250 sites which are associated with 2 kinases"

### 3 Extended Results

Table S2: All AP scores of all pLMs on the 3-NN Model with kinase additional information.

| **Embedding** | **AP** | **+ Family** | **+ Group** | **+ EC** | **+ Family, Group, EC** |
|---|---|---|---|---|---|
| OneHotEnc | 0.0897 | 0.0901 | 0.0868 | 0.0864 | 0.0917 |
| Blosum62 | 0.0897 | 0.0897 | 0.0901 | 0.0901 | 0.0897 |
| NLF | 0.0902 | 0.0903 | 0.0907 | 0.091 | 0.0913 |
| ProtVec | 0.0808 | 0.0963 | 0.1123 | 0.0993 | **0.1230** |
| ProtBERT (avg) | 0.0540 | 0.0871 | 0.0767 | 0.0722 | 0.0985 |
| ProtBERT (cls) | 0.0855 | 0.0867 | 0.0930 | 0.0904 | 0.0968 |
| ProteinBERT | 0.1168 | 0.1182 | 0.1182 | 0.1180 | 0.1227 |
| ProtT5-XL | 0.1172 | 0.1164 | 0.1170 | 0.1172 | 0.1240 |
| Esm1B (avg) | 0.1122 | 0.1107 | 0.1112 | 0.1105 | 0.1132 |
| Esm1B (cls) | 0.1119 | 0.1101 | 0.1110 | 0.1110 | 0.1134 |
| Esm1v (avg) | 0.1114 | 0.1114 | 0.1110 | 0.1112 | 0.1111 |
| Esm1v (cls) | 0.1121 | 0.1125 | 0.1116 | 0.1131 | 0.1129 |
| Esm2 (avg) | 0.0957 | 0.0981 | 0.0977 | 0.1015 | 0.1022 |
| Esm2 (cls) | 0.0982 | 0.1024 | 0.1021 | 0.1024 | 0.1095 |
| DistilProtBERT (avg) | 0.0811 | 0.0961 | 0.0977 | 0.0967 | 0.1085 |
| DistilProtBERT (cls) | 0.1021 | 0.1046 | 0.1084 | 0.1101 | 0.1139 |
| ProtGPT2 | 0.1054 | 0.1061 | 0.1068 | 0.1094 | 0.1111 |
| Ankh-Large | 0.1106 | 0.1077 | 0.1119 | 0.1055 | 0.1137 |
| ProtAlbert (avg) | 0.1004 | 0.1055 | 0.1072 | 0.1041 | 0.1124 |
| ProtAlbert (cls) | 0.0915 | 0.0993 | 0.0958 | 0.0982 | 0.1026 |
| SaProt (avg) | 0.0973 | 0.1109 | 0.1098 | 0.1066 | 0.1209 |
| SaProt (cls) | 0.1056 | 0.1100 | 0.1146 | 0.1107 | 0.1204 |
| TAPE | **0.1200** | **0.1217** | **0.1211** | **0.1249** | **0.1230** |
| ISM2 (avg) | 0.0850 | 0.0878 | 0.0904 | 0.0890 | 0.0965 |
| ISM2 (cls) | 0.0791 | 0.0838 | 0.0813 | 0.0828 | 0.0918 |
| DPLM (avg) | 0.1000 | 0.1006 | 0.0987 | 0.1027 | 0.1034 |
| DPLM (cls) | 0.1034 | 0.1040 | 0.1037 | 0.1051 | 0.1054 |
| AMPLIFY (avg) | 0.0566 | 0.0598 | 0.0604 | 0.0614 | 0.0739 |
| AMPLIFY (cls) | 0.0873 | 0.0870 | 0.0870 | 0.0870 | 0.0871 |
| ESM3 (avg) | 0.1057 | 0.0932 | 0.1049 | 0.1012 | 0.1088 |
| ESM3 (cls) | 0.0896 | 0.0933 | 0.0950 | 0.0901 | 0.1009 |
| ESMC (avg) | 0.0860 | 0.0936 | 0.1052 | 0.0992 | 0.1127 |
| ESMC (cls) | 0.0954 | 0.0942 | 0.1050 | 0.0999 | 0.1090 |
| PTM-Mamba (avg)* | 0.0998 | 0.0946 | 0.0954 | 0.0973 | 0.1007 |
| PTM-Mamba (phosphosite)** | 0.0999 | 0.0939 | 0.0965 | 0.0960 | 0.1004 |

* PTM-Mamba models utilize ESM2 embeddings for kinases.
** Since PTM-Mamba lacks a CLS token and includes special tokens for the phosphorylated residue, we used the embedding of that residue instead.

Table S3: All AP scores of all pLMs on the BZSM Model with kinase additional information.

| Embedding | AP | + Family | + Group | + EC | + Family, Group, EC |
|---|---|---|---|---|---|
| OneHotEnc | 0.0634 | 0.1107 | 0.0832 | 0.0802 | 0.1098 |
| Blosum62 | 0.0327 | 0.0318 | 0.0310 | 0.0337 | 0.0323 |
| NLF | 0.0419 | 0.0391 | 0.0425 | 0.0400 | 0.0426 |
| ProtVec | 0.0959 | 0.1262 | 0.1129 | 0.1214 | 0.1354 |
| ProtBERT (avg) | 0.1044 | 0.1053 | 0.0982 | 0.1017 | 0.1018 |
| ProtBERT (cls) | 0.0842 | 0.1170 | 0.1077 | 0.1132 | 0.1273 |
| ProteinBERT | 0.1236 | 0.1506 | 0.1215 | 0.1367 | 0.1359 |
| ProtT5-XL | 0.1552 | 0.1701 | 0.1531 | 0.1674 | 0.1731 |
| Esm1B (avg) | 0.1512 | 0.1523 | 0.1553 | 0.1546 | 0.1554 |
| Esm1B (cls) | 0.1631 | **0.1740** | **0.1688** | **0.1680** | 0.1769 |
| Esm1v (avg) | 0.1404 | 0.1494 | 0.1426 | 0.1395 | 0.1430 |
| Esm1v (cls) | **0.1640** | 0.1737 | 0.1653 | 0.1652 | 0.1734 |
| Esm2 (avg) | 0.1391 | 0.1588 | 0.1453 | 0.1496 | 0.1638 |
| Esm2 (cls) | 0.1238 | 0.1443 | 0.1358 | 0.1433 | 0.1518 |
| DistilProtBERT (avg) | 0.1269 | 0.1380 | 0.1265 | 0.1356 | 0.1385 |
| DistilProtBERT (cls) | 0.1167 | 0.1360 | 0.1292 | 0.1287 | 0.1441 |
| ProtGPT2 | 0.1333 | 0.1476 | 0.1412 | 0.1419 | 0.1557 |
| Ankh-Large | 0.0840 | 0.1417 | 0.1135 | 0.1178 | 0.1594 |
| ProtAlbert (avg) | 0.0831 | 0.0964 | 0.0854 | 0.0909 | 0.0845 |
| ProtAlbert (cls) | 0.1281 | 0.1269 | 0.1276 | 0.1285 | 0.1372 |
| SaProt (avg) | 0.1466 | 0.1684 | 0.1529 | 0.1490 | 0.1667 |
| SaProt (cls) | 0.1292 | 0.1696 | 0.1424 | 0.1434 | **0.1800** |
| TAPE | 0.1237 | 0.1379 | 0.1333 | 0.1310 | 0.1455 |
| ISM2 (avg) | 0.1124 | 0.1268 | 0.1225 | 0.1193 | 0.1279 |
| ISM2 (cls) | 0.1200 | 0.1275 | 0.1260 | 0.1333 | 0.1374 |
| DPLM (avg) | 0.1299 | 0.1427 | 0.1318 | 0.1368 | 0.1420 |
| DPLM (cls) | 0.1156 | 0.1169 | 0.1145 | 0.1139 | 0.1170 |
| AMPLIFY (avg) | 0.0896 | 0.0968 | 0.0944 | 0.0969 | 0.1066 |
| AMPLIFY (cls) | 0.0969 | 0.0894 | 0.0918 | 0.0928 | 0.0956 |
| ESM3 (avg) | 0.0880 | 0.1544 | 0.1122 | 0.1263 | 0.1585 |
| ESM3 (cls) | 0.0881 | 0.1484 | 0.1220 | 0.1238 | 0.1611 |
| ESMC (avg) | 0.0945 | 0.1502 | 0.1211 | 0.1254 | 0.1615 |
| ESMC (cls) | 0.0866 | 0.1672 | 0.1136 | 0.1401 | 0.1754 |
| PTM-Mamba (avg)* | 0.1092 | 0.1386 | 0.1163 | 0.1230 | 0.1441 |
| PTM-Mamba (phosphosite)** | 0.1218 | 0.1432 | 0.1292 | 0.1346 | 0.1471 |

* PTM-Mamba models utilize ESM2 embeddings for kinases.

** Since PTM-Mamba lacks a CLS token and includes special tokens for the phosphorylated residue, we used the embedding of that residue instead.

Table S4: The mean macro accuracy scores (Acc@k, k=1,3,5) at multiple levels (family, group, phosphosite) for the two best-pLMs, ESM1B (Family + Group + EC) and SaProt (Family + Group + EC), on four random DARKIN splits for the BZSM.

| **Split** | **Embedding** | **Family** | | | **Group** | | |
|---|---|---|---|---|---|---|---|
| | | **Acc@1** | **Acc@3** | **Acc@5** | **Acc@1** | **Acc@3** | **Acc@5** |
| Split 1 | ESM1B (cls) | 0.1404 | 0.3046 | 0.4397 | 0.2739 | **0.4882** | **0.5977** |
| | SaProt (cls) | **0.1526** | **0.3640** | **0.4790** | **0.2886** | 0.4877 | 0.5883 |
| Split 2 | ESM1B (cls) | 0.1398 | 0.2947 | 0.4254 | **0.3150** | **0.4781** | **0.5858** |
| | SaProt (cls) | **0.1481** | **0.3313** | **0.4550** | 0.3067 | 0.4624 | 0.5788 |
| Split 3 | ESM1B (cls) | 0.1722 | 0.3134 | 0.4430 | 0.3560 | 0.5234 | 0.6139 |
| | SaProt (cls) | **0.1776** | **0.3713** | **0.4812** | **0.3579** | **0.5363** | **0.6244** |
| Split 4 | ESM1B (cls) | 0.1664 | 0.3523 | 0.4731 | **0.3095** | 0.5105 | 0.6124 |
| | SaProt (cls) | **0.2043** | **0.4098** | **0.5107** | 0.2992 | **0.5227** | **0.6216** |

### 4 Protein Language Models' Sources

Table S4: Protein language models' versions used in the experiments.

| PLM | Model/Github Link |
|---|---|
| TAPE | github.com/songlab-cal/tape |
| ProtBERT | Rostlab/prot_bert |
| ProtALBERT | Rostlab/prot_albert |
| ProtT5-XL | Rostlab/prot_t5_xl_uniref50 |
| ESM1B | facebook/ESM1B_t33_650M_UR50S |
| ESM1v | facebook/esm1v_t33_650M_UR90S_1 |
| ESM2 | facebook/esm2_t33_650M_UR50D |
| ProteinBERT | github.com/nadavbra/protein_bert |
| ProtGPT2 | nferruz/ProtGPT2 |
| DistilProtBERT | yarongef/DistilProtBert |
| Ankh | ElnaggarLab/ankh-large |
| SaProt | westlake-repl/SaProt_650M_PDB |
| ESM3 | github.com/evolutionaryscale/esm(1.4B) |
| ESMC | github.com/evolutionaryscale/esm(600M) |
| ISM2 | jozhang97/ism_t33_650M_uc30pdb |
| DPLM | github.com/bytedance/dplm/tree/representationlearning |
| AMPLIFY | github.com/chandar-lab/AMPLIFY(350M) |
| PTM-Mamba | github.com/programmablebio/ptm-mamba |

### 5 Fine Tuning with Auxilary Kinase and Phosphosite Prediction Tasks

**i) Fine-tuning based on Phosphorylation Prediction Task:**

In this task, given a 15-mer peptide sequence surrounding the potential phosphosite (Ser, Thr, or Tyr), the model learns to predict whether the site can be phosphorylated or not. We retrieved 366.028 experimentally phosphorylated site information from PhosphositePlus (March 18, 2024). This constitutes the positive class. For the negative-labeled peptide sequences, we followed a common approach. [Wang et al., 2017] For every positively labeled phosphosite, we retrieve the protein sequence it is in and pick another Ser, Thr, or Tyr within the same protein sequence that is not in Phosphositeplus (not labeled as a phosphosite). Centered on this site, the 15-mer constitutes a negatively labeled phosphosite. This procedure yielded 730,149 distinct 15-mer peptides. To fine-tune ESM1 b, we split the training data into train-validation-test sets by 70-15-15% ratio. We employed validation data to optimize the pLM's phosphorylation prediction performance during fine-tuning. We used AdamW as the optimizer, the learning rate started from 5e-5, the batch size was set to 8, and we fine-tuned for 3 epochs.

We checked if the model learns to classify the phosphorylation site correctly. The model achieves 0.94 accuracy on the test data. Note that the dataset is balanced in terms of positive and negative labeled examples.

##### ii) Fine-tuning Kinase Embeddings Based on Kinase-Group Classification:

To arrive at a kinase-aware PLM, we fine-tuned the kinase embeddings on the kinase group classification task. We used 392 human kinase domains' sequences and the kinases group information. There are a total of 10 groups. We split the dataset into train-validation-test sets by a 70-15-15% ratio. We used the AdamW optimizer. The learning rate started from 5e-5 and determined the batch size as 8. We fine-tuned ESM1b three epochs.

The Table S5 presents the results of the BZSM results. The first rows in the tables show the results of cases if both representations are retrieved from the general-purpose ESM1b, which we used earlier in the paper. The remaining rows represent the cases if we use a fine-tuned model representation. The results show that fine-tuning the models with kinase data actually did not produce good representations. Here, the main reason probably is the small number of kinases.

Table S5: The experiments are conducted on BZSM by using fine-tuned embeddings. The results in the first row are the reference result, from which both kinase and phosphosite representations are obtained from the general-purpose ESM1b. The rest of the experiments are performed by changing peptide representations to see the effect of fine-tuned embeddings.

| Phosphosite Representation | Kinase Representation | Kinase Features | AP |
|---|---|---|---|
| ESM1b | ESM1b | Seq,Family,Group,EC | 0.1769 |
| Fine-tuned ESM1b on Phosphorylation Prediction | ESM1b | Seq,Family,Group,EC | 0.1796 |
| Fine-tuned ESM1b on Phosphorylation Prediction | Fine-tuned ESM1b on Group Prediction | Seq,Family,EC | 0.1088 |
| Fine-tuned ESM1b on Phosphorylation Prediction | Fine-tuned ESM1b on Group Prediction | Seq,Family,Group,EC | 0.1074 |

##### iii) Fine-tuning Kinase Embeddings with Contrastive Learning Based on Functional and Family Similarities

We have further detailed our fine-tuning approach for kinase embeddings, which leverages biologically-informed contrastive learning based on kinase family/group relationships and amino acid target specificity. We prepared a dataset by creating positive and negative pairs based on family and group information. Kinases belonging to the same family and phosphorylating the same type of amino acid (S, T, or Y) were determined as positive pairs, while kinases belonging to different families and phosphorylating the same type of amino acid as the positive kinases were determined as negative pairs. Finally, we obtained 79,000 positive and negative pairs for kinase groups and 83,000 positive and negative pairs for kinase families. We fine-tuned ESM-1b (650M parameters) 6 epochs using LoRA with 4-bit quantization. We used AdamW as the optimizer, the learning rate started from 1e-4, and the batch size was set to 128. The results are shown in Table S6.

Fine-tuning ESM1b with positive and negative kinase family/group pairs yielded APs of 0.1792–0.1785,

Table S6: Experiments to see the effect of fine-tuning models learning family/group specificity.

| **Phosphosite Representation** | **Kinase Representation** | **Kinase Features** | **AP** |
|---|---|---|---|
| ESM1b | ESM1b | Seq,Family,Group,EC | 0.1769 |
| ESM1b | Fine-tuned ESM1b on Kinase Group Pairs | Seq,Family,Group,EC | 0.1785 |
| ESM1b | Fine-tuned ESM1b on Kinase Family Pairs | Seq,Family,Group, EC | 0.1792 |

a very modest lift over the case using the general purpose ESM1b. This demonstrates that simple family/group-based positive/negative sampling provides only a marginal benefit. The small gains may stem from intra-family/group substrate heterogeneity.
